# Gene nucleotide composition accurately predicts expression and is linked to topological chromatin domains

**DOI:** 10.1101/117499

**Authors:** Chloé Bessière, May Taha, Florent Petitprez, Jimmy Vandel, Jean-Michel Marin, Laurent Bréhélin, Sophie Lèbre, Charles-Henri Lecellier

## Abstract

Gene expression is orchestrated by distinct regulatory regions (e.g. promoters, enhancers, UTRs) to ensure a wide variety of cell types and functions. A challenge is to identify which regulatory regions are active, what are their associated features and how they work together in each cell type. Several approaches have tackled this problem by modeling gene expression based on epigenetic marks (e.g. ChIP-seq, methylation, DNase hypersensitivity), with the ultimate goal of identifying driving genomic regions and mutations that are clinically relevant in particular in precision medicine. However, these models rely on experimental data, which are limited to specific samples (even often to cell lines) and cannot be generated for all regulators and all patients. In addition, we show here that, although these approaches are accurate in predicting gene expression, their biological interpretation can be misleading. Finally these methods are not designed to capture potential regulation instructions present at the sequence level, before the binding of regulators or the opening of the chromatin. We develop here a method for predicting mRNA levels based solely on sequence features collected from distinct regulatory regions, which is as accurate as methods based on experimental data. Our approach confirms the importance of nucleotide composition in predicting gene expression and ranks regulatory regions according to their contribution. It also unveils strong influence of gene body sequence, in particular introns. We further provide evidence that the contribution of nucleotide content can be linked to co-regulations associated with genome 3D architecture and to associations of genes within topologically associated domains.

## INTRODUCTION

The diversity of cell types and cellular functions is defined by specific patterns of gene expression. The regulation of gene expression involves a plethora of DNA/RNA-binding proteins that bind specific motifs present in various DNA/RNA regulatory regions. At the DNA level, transcription factors (TFs) typically bind 6-8bp-long motifs present in promoter regions, which are close to transcription start site (TSS). TFs can also bind enhancer regions, which are distal to TSSs and often interspersed along considerable physical distance through the genome [1]. The current view is that DNA looping mediated by specific proteins and RNAs places enhancers in close proximity with target gene promoters (for review [2, 3, 4, 5]). High-resolution chromatin conformation capture (Hi-C) technology identified contiguous genomic regions with high contact frequencies, referred to as topologically associated domains (TADs) [6]. Within a TAD, enhancers can work with many promoters and, on the other hand, promoters can contact more than one enhancer [7, 5].

At the RNA level, RNA-binding proteins (RBPs) can co- or post-transcriptionally regulate the fate of RNAs. The vast majority of RBPs appears to bind target sequences in single-strand RNA, and none absolutely requires a specific RNA secondary structure [8, 9, 10]. Some RBPs guided by small noncoding RNAs i.e. microRNAs (miRNAs) can also recognize specific sequences in their RNA targets leading eventually to RNA degradation [11]. The miRNAs are thought to fine-tune the RNA levels but their regulatory power appears overshadowed by TFs [12]. The RNA regions involved in post-transcriptional regulations mostly correspond to the 5’ and 3’ untranslated regions (5’UTR and 3’UTR) [8, 13, 14]. The coding DNA sequence (CDS) also plays key role in posttranscriptional [15, 16, 17, 18] as well as transcriptional [19] regulations. Note that the situation is obviously different in the case of non coding RNAs, which are not considered in this study. Intronic sequences have also been reported to affect gene regulation in many ways, from transcription to RNA stability, that are distinct from splicing [20, 21].

Several large-scale data derived from high-throughput experiments (such as ChIP-seq [22], SELEX-seq [23], RNAcompete [24]) can be used to highlight TF/RBP binding preferences and build Position Weight Matrixes (PWMs) [25]. The human genome is thought to encode ~2,000 TFs [26] and >1,500 RBPs [14]. It follows that gene regulation is achieved primarily by allowing the proper combination to occur i.e. enabling cell-and/or function-specific regulators (TFs or RBPs) to bind the proper sequences in the appropriate regulatory regions. In that context, epigenetics clearly plays a central role as it influences the binding of the regulators and ultimately gene expression [27]. Provided the variety of regulatory mechanisms, deciphering their combination requires mathematical/computational methods able to consider all possible combinations.

Several methods have recently been proposed to tackle this problem [28, 12, 29, 30]. Although these models appear very efficient in predicting gene expression and identifying key regulators, they mostly rely on experimental data (ChIP-seq, methylation, DNase hypersensitivity), which are limited to specific samples (often to cell lines) and which cannot be generated for all TFs/RBPs and all cell types. These technological features impede from using this type of approaches in a clinical context in particular in precision medicine. In addition, we show here that, although these approaches are accurate, their biological interpretation can be misleading. Finally these methods are not designed to capture regulation instructions that may lie at the sequence-level before the binding of regulators or the opening of the chromatin. There is indeed a growing body of evidence suggesting that the DNA sequence *per se* contains information able to shape the epigenome and explain gene expression [31, 32, 33, 34, 35]. Several studies have shown that sequence variations affect histone modifications [32, 33, 34]. Specific DNA motifs can be associated with specific epigenetic marks and the presence of these motifs can predict the epigenome in a given cell type [35]. Quante and Bird proposed that proteins able to ”read” domains of relatively uniform DNA base composition may modulate the epigenome and ultimately gene expression [31]. In that view, modeling gene expression using only DNA sequences and a set of predefined DNA/RNA features (without considering experimental data others than expression data) would be feasible. In line with this proposal, Raghava and Han developed a Support Vector Machine (SVM)-based method to predict gene expression from amino acid and dipeptide composition in *Saccharomyces cerevisiae* [16].

Here, we built a global regression model per sample to predict the expression of the different genes using their nucleotide compositions as predictive variables. The idea beyond our approach is that the selected variables (defining the model) are specific to each sample. Hence the expression of a given gene may be predicted by different variables in different samples. This approach was tested on several RNA-sequencing data from The Cancer Genome Atlas (TCGA) and showed accuracy similar to that of methods based on experimental data. We thus confirmed the importance of nucleotide composition in predicting gene expression. Moreover, the gene body (introns, CDS and UTRs), as opposed to sequences located upstream (promoter) or downstream, had the most significant contribution in our model. We further provided evidence that the contribution of nucleotide composition in predicting gene expression is linked to co-regulations associated with genome architecture and TADs.

## MATERIAL AND METHODS

### Datasets, sequences and online resources

RNA-seq V2 level 3 processed data were downloaded from the TCGA Data Portal. Our training data set contained 241 samples randomly chosen from 12 different cancers (20 cancerous samples for each cancer except 21 for LAML, see Supplementary Table S7). Isoform expression data (.rsem.isoforms.normalized_results files) were downloaded from the Broad TCGA GDAC (http://gdac.broadinstitute.org) using firehose_get. We collected data for 73599 isoforms in 225 samples of the 241 initially considered. All the genes and isoforms not detected (no read) in any of the considered samples were removed from the analyses. Expression data were log transformed.

All sequences were mapped to the hg38 human genome and the UCSC liftover tool was used when necessary. Gene TSS positions were extracted from GENCODEv24. UTR and CDS coordinates were extra@cted from ENSEMBL Biomart. To assign only one 5UTR sequence to one gene, we merged all annotated 5UTRs associated with the gene of interest using Bedtools merge [36] and further concatenated all sequences. The same procedure was used for 3UTRs and CDSs. Intron sequences are GENCODEv24 genes to which 5UTR, 3UTR and CDS sequences described above were substracted using Bedtools substract [36]. These sequences therefore corresponded to constitutive introns. The intron sequences were concatenated per gene. The downstream flanking region (DFR) was defined as the region spanning 1kb after GENCODE v24 gene end. Fasta files were generated using UCSC Table Browser or Bedtools getfasta [36].

TCGA isoform TSSs were retrieved from https://webshare.bioinf.unc.edu/public/mRNAseq_TCGA/unc_hg19.bed and converted into hg38 coordinates with UCSC liftover. For other regulatory regions associated to transcript isoforms (UTRs, CDS, introns and DFR), we used GENCODE v24 annotations.

### Nucleotide composition

The nucleotide (n=4) and dinucleotide (n=16) percentages were computed from the different regulatory sequences where:

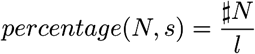

is the percentage of nucleotide *N* in the regulatory sequence *s*, with *N* in {*A, C, G, T*} and *l* the length of sequence *s*, and

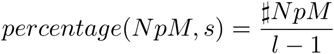

is the *NpM* dinucleotide percentage in the regulatory sequence *s*, with *N* and *M* in {*A, C, G, T*} and *l* the length of sequence *s*.

### Motif scores

Motif scores in core promoters were computed using the method explained in [25] and Position Weight Matrix (PWM) available in JASPAR CORE 2016 database [37]. Let *w* be a motif and *s* a nucleic acid sequence. For all nucleotide *N* in {*A, C, G, T*}, we denoted by *P*(*N*|*w_j_*) the probability of nucleotide *N* in position *j* of motif *w* obtained from the PWM, and by *P*(*N*) the prior probability of nucleotide *N* in all sequences.

The score of motif *w* at position *i* of sequence *s* is computed as follows:

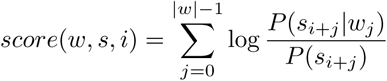

with |*w*| the length of motif *w*, *s_i_*_+*j*_ the nucleotide at position *i* + *j* in sequence *s*, The score of motif *w* for sequence *s* is computed as the maximal score that can be achieved at any position of *s*, i.e.:

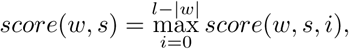

with *l* the length of sequence *s*.

### DNAshape scores

DNA shape scores were computed using DNAshapeR [38].

### Enhancers

The coordinates of the enhancers mapped by FANTOM on the hg19 assembly [7] were converted into hg38 using UCSC liftover and further intersected with the different regulatory regions. We computed the density of enhancers per regulatory region (*R*) by dividing the sum, for all genes, of the intersection length of enhancers with gene *i* (*L_enhi_*) by the sum of the lengths of this regulatory region for all genes:

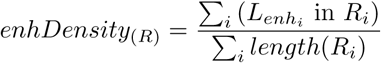

### Expression quantitative trait loci

The v6p GTex *cis*-eQTLs were downloaded from the GTex Portal (http://www.gtexportal.org/home/). The hg19 *cis*-eQTL coordinates were converted into hg38 using UCSC liftover and further intersected with the different regulatory regions. We restricted our analyses to *cis*-eQTLs impacting their own host gene. We computed the density of *cis*-eQTL per regulatory region (*R*) by dividing the sum, for all genes, of the number of *cis*-eQTLs of gene *i* (*eQTLs_i_*) located in the considered region for gene *i* (*R_i_*) by the sum of the lengths of this regulatory region for all genes:

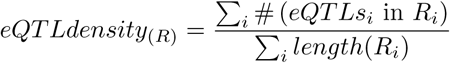

### Methylation

Illumina Infinium Human DNA Methylation 450 level 3 data were downloaded from the Broad TCGA GDAC (http://gdac.broadinstitute.org) using firehose_get. The coordinates of the methylation sites (hg18) were converted into hg38 using the UCSC liftover and further intersected with that of the core promoters (hg38). For each gene, we computed the median of the beta values of the methylation sites present in the core promoter and further calculated the median of these values in 21 LAML and 17 READ samples with both RNA-seq and methylation data. We compared the overall methylation status of the core promoters in LAML and READ using a wilcoxon test.

### Functional enrichment

Gene functional enrichments were computed using the database for annotation, visualization and integrated discovery (DAVID) [39].

### Linear regression with *ℓ*_1_-norm penalty (Lasso)

We performed estimation of the linear regression model (1) via the lasso [40]. Given a linear regression with standardized predictors and centered response values, the lasso solves the *ℓ*_1_-penalized regression problem of finding the vector coefficient *β* = {*β_ί_*} in order to minimize

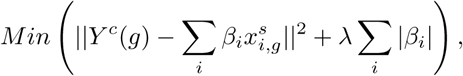

where *Y*^c^(*g*) is the centered gene expression for all gene *g*, 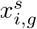 is the standardized DNA feature *i* for gene *g* and Σ_*i*_ |*β_i_*| is the *ℓ*_1_-norm of the vector coefficient *β*. Parameter λ is the tuning parameter chosen by 10 fold cross validation. The highest value of λ, the less variables selected. This is equivalent to minimizing the sum of squares with a constraint of the form Σ_*i*_ |*β_i_*| ≤ *s*. Gene expression predictions are computed using coefficient *β* estimated with the value of λ that minimizes the mean square error. Lasso inference was performed using the function cv.glmnet from the R package glmnet [41].

### Stable variable selection

Consistently selected variables were identified using the two functions stabpath and stabsel from the R package C060 for glmnet models [42]. In the first step, for each sample, the lasso inference is repeated 500 times such that, for each iteration, only 50% of the genes is used (uniformly sampled) and each predictive variable is reweighted by a random weight (uniformly sampled in [0.5; 1]). In the second step, variables are considered as stable if they are selected in more than 70% of the iterations using the method proposed in [42] to set the value of λ.

### Regression trees

Regression trees were implemented with the rpart package in R [41]. In order to avoid over-fitting, trees were pruned based on a criterion chosen by cross validation to minimize mean square error. The minimum number of genes was set to 100 genes per leaf.

### TAD enrichment

We considered TADs mapped in IMR90 cells [6] containing more than 10 genes (373 out of 2243 TADs with average number of genes = 14). The largest TAD had 76 associated genes. First, for each TAD and for each region considered, the percentage of each nucleotide and dinucleotide associated to the embedded genes were compared to that of all other genes using a Kolmogorov-Smirnov test. Correction for multiple tests was applied using the False Discovery Rate (FDR) < 0.05 [43] and the R function p.adjust [41]. Second, for each of the 967 groups of genes (identified by the regression trees, with mean error < mean error of the 1st quartile), the over-representation of each TAD within each group was tested using the R hypergeometric test function phyper [41]. Correction for multiple tests was applied using FDR< 0.05 [43].

### Cancer acronyms

ovarian serous cystadenocarcinoma, OV; bladder urothelial carcinoma, BLCA; breast invasive carcinoma, BRCA; colon adenocarcinoma, COAD; lymphoid neoplasm diffuse large B-cell lymphoma, DLBC; acute myeloid leukemia, LAML; brain lower grade glioma, LGG; liver hepatocellular carcinoma, LIHC; lung adenocarcinoma, LUAD; pancreatic adenocarcinoma, PAAD; prostate adenocarcinoma, PRAD; rectum adenocarcinoma, READ.

### Data availability

The matrices of predicted variables (log transformed RNA seq data) and predictive variables (nucleotide and dinucleotide percentages, motifs and DNA shape scores computed for all genes as described above) as well as the TCGA barcodes of the 241 samples used in our study have been made available at http://www.univ–montp3.fr/miap/~lebre/IBCRegulatoryGenomics.

## RESULTS

### Mathematical approach to predict gene expression

We built a global linear regression model to predict the expression of genes using DNA/RNA features associated with their regulatory regions (e.g. nucleotide composition, TF motifs, DNA shapes):

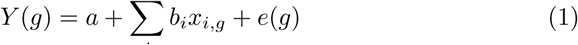

where *Y*(*g*) is the expression of gene *g, x_i_,_g_* is feature *i* for gene *g, e*(*g*) is the residual error associated with gene *g, a* is the intercept and *b_i_* is the regression coefficient associated with feature *i*.

The advantage of this approach is that it allows to unveil, into a single model, the most important regulatory features responsible for the observed gene expression. The relative contribution of each feature can thus be easily assessed. It is important to note that the model is specific to each sample. Hence the expression of a given gene may be predicted by different variables depending on the sample. Our computational approach was based on two steps. First, a linear regression model (1) was trained with a lasso penalty [40] to select sequence features relevant for predicting gene expression. Second, the performances of our model was evaluated by computing the mean square of the residual errors, and the correlation between the predicted and the observed expression for all genes. Our approach was applied to a set of RNA sequencing data from TCGA. We randomly selected 241 gene expression data from 12 cancer types (see http://www.univ–montp3.fr/miap/~lebre/IBCRegulatoryGenomics for the barcode list). For each dataset (i.e sample), a regression model was learned and evaluated. See Materials and Methods for a complete description of the data, the construction of the predictor variables and the inference procedure.

### Contribution of the promoter nucleotide composition

We first evaluated the contribution of promoters, which are one of the most important regulatory sequences implicated in gene regulation [44]. We extracted DNA sequences encompassing ±2000 bases around all GENCODE v24 TSSs and looked at the percentage of dinucleotides along the sequences (Supplementary Figure S1). Based on these distributions, we segmented the promoter into three distinct regions: −2000/−500 (referred here to as distal upstream promoter, DU), −500/+500 (thereafter called core promoter though longer than the core promoter traditionally considered) and +500/+2000 (distal downstream promoter, DD)(Figure 1). We computed the nucleotide (n=4) and dinucleotide (n=16) relative frequencies in the three distinct regions of each gene. For each sample, we trained one model using the 20 nucleotide/dinucleotide relative frequencies from each promoter segment separately, and from each combination of promoter segments. We observed that the core promoter had the strongest contribution compared to DU and DD (Figure 2(a)). Considering promoter as one unique sequence spanning −2000/+2000 around TSS achieved lower model accuracy than combining different promoter segments (Figure 2(a)). The highest accuracy was obtained combining all three promoter segments Figure 2(a)).

**Figure 1.**
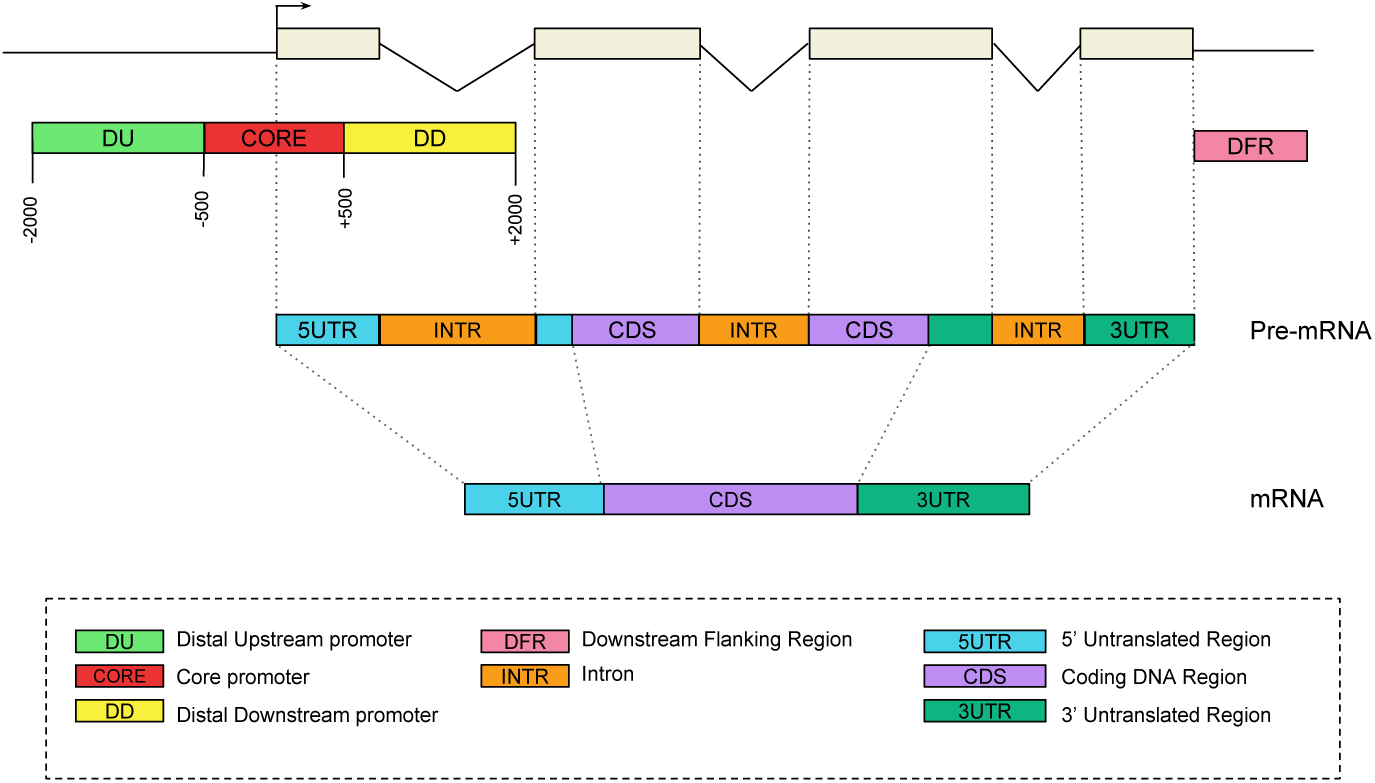
Genomic regions considered for gene expression prediction. An illustrative transcript is shown as example.

**Figure 2.**
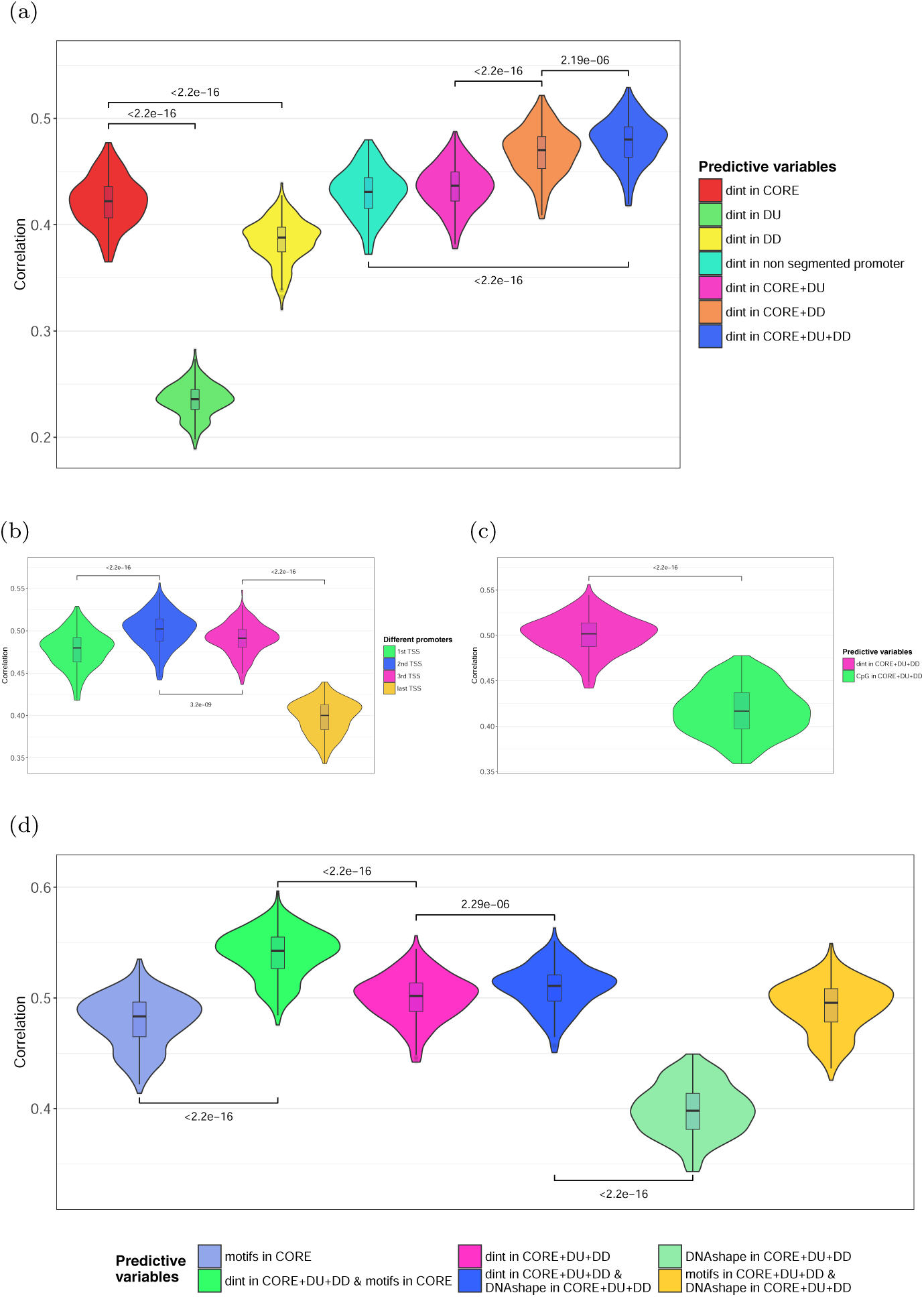
**(a) Contribution of the promoter segments.** The model was built using 20 variables corresponding to the nucleotide (4) and dinucleotide (16) percentages computed in the CORE promoter (red), DU (green) or DD (yellow). These variables were then added in different combinations: CORE+DU (pink, 40 variables); CORE+DD (orange, 40 variables); CORE+DU+DD (light blue, 60 variables). Promoter segments were centered around the first most upstream TSS. The model was also built on 20 variables corresponding to the nucleotide and dinucleotide compositions of the non segmented promoters (-2000/+2000 around the first most upstream TSS)(light blue). All different models were fitted on 19393 genes for each of the 241 samples considered. The prediction accuracy was evaluated in each sample by evaluating the Spearman correlation coefficients between observed and predicted gene expressions. The correlations obtained in all samples are shown as violin plots. **(b) Prediction accuracy comparing alternative TSSs.** The model was built using the 60 nucleotide/dinucleotide percentages computed in the 3 promoter segments (CORE+DU+DD) centered around 1st, 2nd, 3rd and last TSSs. **(c) Contribution of CpG.** The model was built using the 60 nucleotide/dinucleotide or only the 3 CpG percentages computed in the 3 promoter segments (CORE+DU+DD) centered around the 2nd TSS. **(d) Contribution of motifs and local DNA shapes**. The model was built using (i) 60 nucleotide/dinucleotide percentages computed in the 3 promoter segments (CORE+DU+DD) (“dint”, pink),(ii) 471 JASPAR2016 PWM scores computed in the CORE segment (“motifs”, light blue) and (iii) the 12 DNA shapes corresponding to the 4 known DNAshapes computed in CORE, DU and DD (“DNAshape”, green). All sequences were centered around the 2nd TSS. These variables were further added in different combinations to build the models indicated: dint+motifs (531 variables, green), dint+DNAshapes (32 variables, dark blue), motifs+DNAshapes (483 variables, light green).

Promoters are often centered around the 5’ most upstream TSS (i.e. gene start). However genes can have multiple transcriptional start sites. The median number of alternative TSSs for the 19,393 genes listed in the TCGA RNA-seq V2 data is 5 and only 2,753 genes harbor a single TSS (Supplementary Figure S2). We therefore evaluated the performance of our model comparing different promoters centered around the first, second, third and last TSS (Figure 2(b)). In the absence of second TSS, we used the first TSS and likewise the second TSS in the absence of a third TSS. The last TSS represents the most downstream TSS in all cases. We found that our model achieved higher predictive accuracy with the promoters centered around the second TSS (Figure 2(b)), in agreement with [28].

We noticed that incorporating the number of TSSs associated with each gene drastically increased the performance of our model (Supplementary Figure S3). Multiplying TSSs may represent a genuine mechanism to control gene expression level. On the other hand this effect may merely be due to the fact that the more a gene is expressed, the more its different isoforms will be detected (and hence more TSSs will be annotated). Because the number of known TSSs results from annotations deduced from experiments, we decided not to include this variable into our final model.

### Contribution of specific features associated with promoters

Provided the importance of CpGs in promoter activity [44], we first compared our model with a model built only on promoter CpG content. We confirmed that CpG content had an important contribution in predicting gene expression (median R = 0.417, Figure 2(c)). However considering other dinucleotides achieved better model performances, indicating that dinucleotides other than CpG contribute to gene regulation. This is in agreement with results obtained by Nguyen *et al.*, who showed that CpG content is insufficient to encode promoter activity and that other features might be involved [45].

We integrated TF motifs considering Position Weight Matrix scores computed in the core promoter and observed a slight but significant increase of the regression performance (median r = 0.543 with motif scores vs. r = 0.502 without motif scores, Figure 2(d)). As three-dimensional local structure of the DNA (DNA shape) improve TF motif predictions [46], we also computed, for each promoter segment, the mean scores of the four DNA shape features provided by DNAshapeR [38] (helix twist, minor groove width, propeller twist, and Roll). Although the difference between models with and without DNA shapes is also significant, the increase in performance is more modest than when including TF motif scores (Figure 2(d)).

Our model suggested that nucleotide composition had a greater contribution in predicting gene expression compared to TF motifs and DNA shapes. This is in agreement with the findings revealing the influence of the nucleotide environment in TFBS recognition [47]. Note however that nucleotide composition, TF motifs and DNA shapes may be redundant variables. Besides, a linear model may not be optimal to efficiently capture the contributions of TF motifs and/or DNA shapes. The highest performance was achieved by combining nucleotide composition with TF motifs (Figure 2(d)). In the following analyses, the model was built on both dinucleotide composition and core promoter TF motifs.

### Comparison with models based on experimental data

The wealth of TF ChIP-seq, epigenetic and expression data has allowed the development of methods aimed at predicting gene expression based on differential binding of TFs and epigenetic marks [28, 12, 29, 30]. We sought to compare our approach, which does not necessitate such cell-specific experimental data, to these methods. We first compared our results to that of Li *et al.* who used a regression approach called RACER to predict gene expression on the basis of experimental data, in particular ChIP-seq data and DNA methylation [12]. Note that, with this model, the contribution of TF regulation in predicting gene expression is higher than that of DNA methylation [12].

We computed the Spearman correlations between expressions observed in the subsets of LAMLs studied in [12] and expressions predicted by our model or by RACER (Figure 3(a)). For the sake of comparison, we used the RACER model built solely on ChIP-seq data, hereafter referred to as ”ChIP-based model”. Overall our model was as accurate as ChIP-based model (median correlation r = 0.529 with our model vs. median r = 0.527 with ChIP-based model (Figure 3(a))). We then controlled the biological information retrieved by the two approaches by randomly permuting, for each gene, the values of the predictive variables (dinucleotide counts/motif scores in our model and ChIP-seq signals in the ChIP-based model). This creates a situation where the links between the combination of predictive variables and expression is broken, while preserving the score distribution associated with each gene. In such situation, a regression model is expected to poorly perform. Surprisingly, the accuracy of ChIP-based model was not affected by the randomization process (median r = 0.517, Figure 3(a)) while that of our model was severely impaired (median r = 0.076, Figure 3(a)). We built another control model using a single predictive variable per gene corresponding to the maximum value of all predictive variables initially considered. Here again the ChIP-based model was not affected by this process (median r = 0.520, Figure 3(a)) while our model failed to accurately predict gene expression with this type of control variable (median r = −0.016, Figure 3(a)).

**Figure 3.**
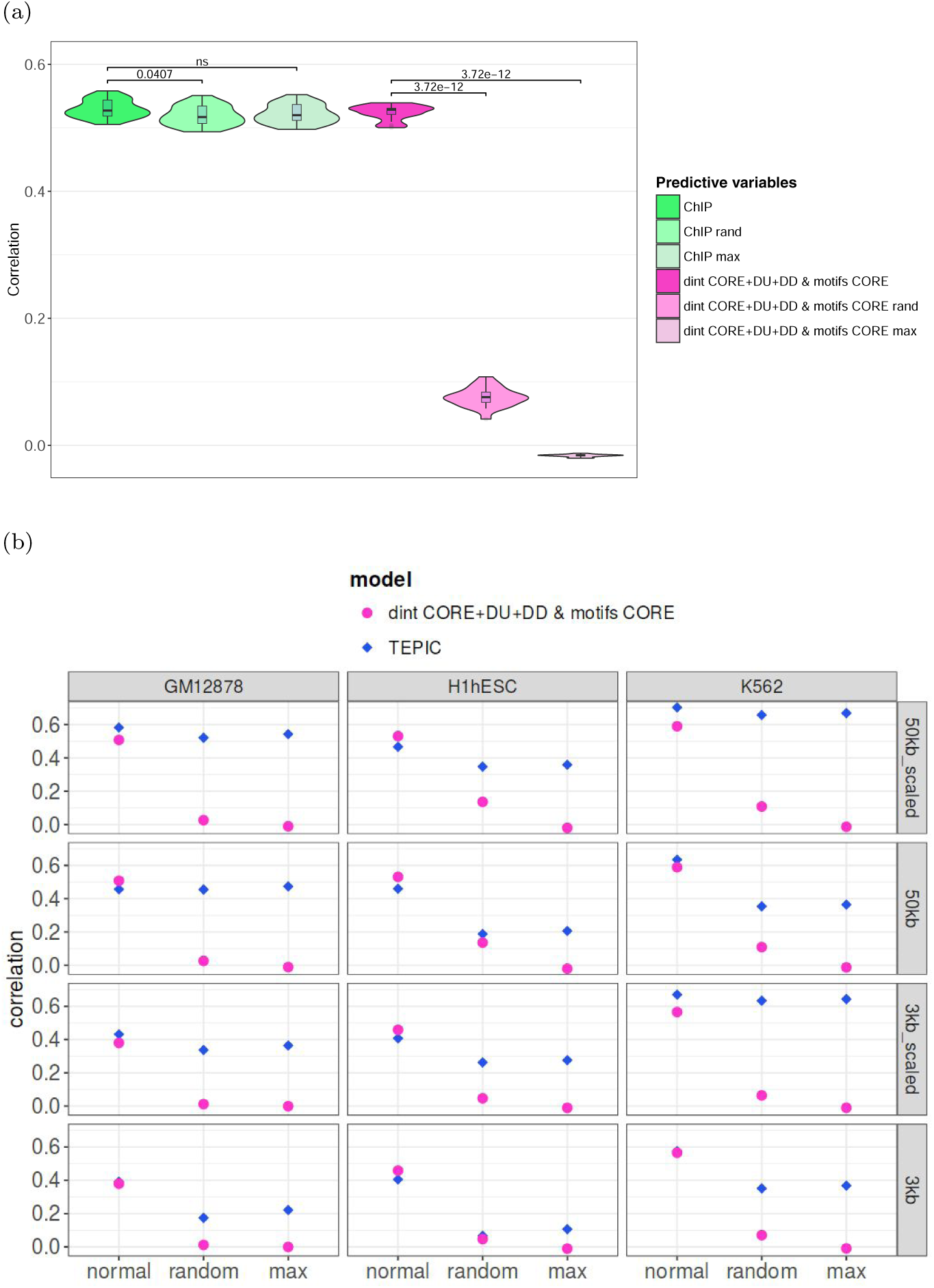
**(a) Comparison with model integrating TF-binding signals.** The model was built using 531 variables corresponding to the 60 nucleotide/dinucleotide percentages and the 471 motif scores computed in the 3 promoter segments (CORE, DU, DD) centered around the 2nd TSS (pink). A model built on ChIP-seq data [12] was used for comparison (green). Both models were fitted on the same gene set (n=16298) for 21 LAML samples. The two models were also built on randomized values of predictive variables (rand) and on the maximum value of all predictive variables (max). **(b) Comparison with model integrating open-chromatin signals.** The linear model was built using the 531 variables (nucleotide/dinucleotide percentages and motif scores in CORE, DU and DD) and the expression data obtained in K562, hESC and GM12878 [30]. TEPIC was built as described in [30], within a 3 kb or a 50 kb window around TSSs. The scaled version of TEPIC incorporates the abundance of open-chromatin peaks in the analyzed sequences. All types of TEPIC models were tested (3kb, 3kb-scaled, 50kb and 50kb-scaled). In each case, our model was built on the set of genes considered by TEPIC. Models were further built on randomized values of predictive variables (rand) and on the maximum value of all predictive variables (max).

ChIP-seq data are probably the best way to measure the activity of a TF because binding of DNA reflects the output of RNA/protein expression as well as any appropriate post-translational modifications and subcellular localizations. However this type of data also reflects chromatin accessibility (i.e. most TFs bind accessible genomic regions) and TFs tend to form clusters on regulatory regions [48]. The binding of one TF in the promoter region is therefore likely accompanied by the binding of others. Hence, rather than inferring the TF combination responsible for gene expression, linear models based of ChIP-seq data may capture the quantity of TFs (i.e. the opening of the chromatin) in the promoter region of each gene, which explains their good accuracy on randomized or maximized variables.

We indeed observed a similar bias in the results obtained by TEPIC [30], a regression method that predicts gene expression from PWM scores and open-chromatin data. Specifically, TEPIC computes a TF-affinity score for each gene and each PWM by summing up the TF affinities in all open-chromatin peaks (DNaseI-seq) within a close (3,000 bp) or large (50,000 bp) window around TSSs. This scoring takes into account the scores of PWMs in the open-chromatin peaks but is also influenced by the number of open-chromatin peaks in the analyzed sequences and the abundance of open-chromatin peaks (“scaled” version). As a result, genes with many open-chromatin peaks tend to get higher TF-affinity scores than genes with low number of open-chromatin peaks. We trained linear models on three cell-lines using either the four TEPIC affinity-scores or our variables and compared the results (Figure 3(b)). As for the ChIP-based models, we observed that our model was approximately as accurate as TEPIC score model. Applying the random permutations on the TEPIC scores did not significantly impact the accuracy of the approach in most cases, especially for the scaled versions (Figure 3(b)). Hence, as for the ChIP-based model, the TEPIC score model seems to mainly capture the level of chromatin opening rather than the TF combinations responsible for gene expression. Conversely, our model solely built on DNA sequence features is not influenced by the chromatin accessibility and thus can yield relevant combinations of explanatory features (see the randomized control in Figures 3(a) and 3(b)). Note that the non-scaled version of TEPIC did show a loss of accuracy for cell-line H1-hESC (as well as a moderate loss for K562, but none for GM12878) when randomizing or maximizing the variables (Figure 3(b)). This result indicates that, although taking the abundance of open-chromatin peaks in the analyzed sequences does increase expression prediction accuracy, it might generate more irrelevant combinations of explanatory features than non-scaled versions.

### Contribution of additional genomic regions

Additional genomic regions were integrated into our model. We first thought to consider enhancer sequences implicated in transcriptional regulation. We used the enhancer mapping made by the FANTOM5 project, which identified 38,554 human enhancers across 808 samples [7]. This mapping uses the CAGE technology, which captures the level of activity for both promoters and enhancers in the same samples. It is then possible to predict the potential target genes of the enhancers by correlating the activity levels of these regulatory regions over hundreds of human samples [7]. However FANTOM5 enhancers are only assigned to 11,359 genes from the TCGA data, which correspond to the most expressed genes across different cancers (Supplementary Figure S4). Provided that the detection of enhancers relies on their activity, it is expected that enhancers are better characterized for the most frequently expressed genes. Because considering enhancers would considerably reduce the number of genes and introduce a strong bias in the performance of our model, we decided not to include these regulatory regions.

Second we analyzed the contribution of regions defined at the RNA level namely 5’UTR, CDS, 3’UTR and introns (Figure 1). For all genes, we extracted all annotated 5’UTRs, 3’UTRs and CDSs, which were further merged and concatenated to a single 5’UTR, a single CDS, and a single 3’UTR per gene. Introns were defined as the remaining sequence (Figure 1). We also tested the potential contribution of the 1kb region located downstream the gene end, called thereafter Downstream Flanking Region (DFR, Figure 1). Our rationale was based on reports showing the presence of transient RNA downstream of polyadenylation sites [49], the potential presence of enhancers [7] and the existence of 5’ to 3’ gene looping [50].

We used a forward selection procedure by adding one region at a time: (i) all regions were tested separately and the region leading to the highest Spearman correlation between observed and predicted expression was selected as the ‘first’ seed region, (ii) each region not already in the model was added separately and the region yielding the best correlation was selected (‘second region’), (iii) the procedure was repeated till all regions were included in the model. The correlations computed at each steps are indicated in Supplementary Table S1. As shown in Figure 4, the nucleotide composition of intronic sequences had the strongest contribution in the accuracy of our model, followed by UTRs (5’ then 3’) and CDS (Figure 4). The nucleotide composition of core promoter moderately increased the prediction accuracy. In contrast the composition of regions flanking core promoter (DU and DD, Figure 1) as well as regions located downstream the end of gene (DFR, Figure 1) did not significantly improve the predictions of our model.

**Figure 4.**
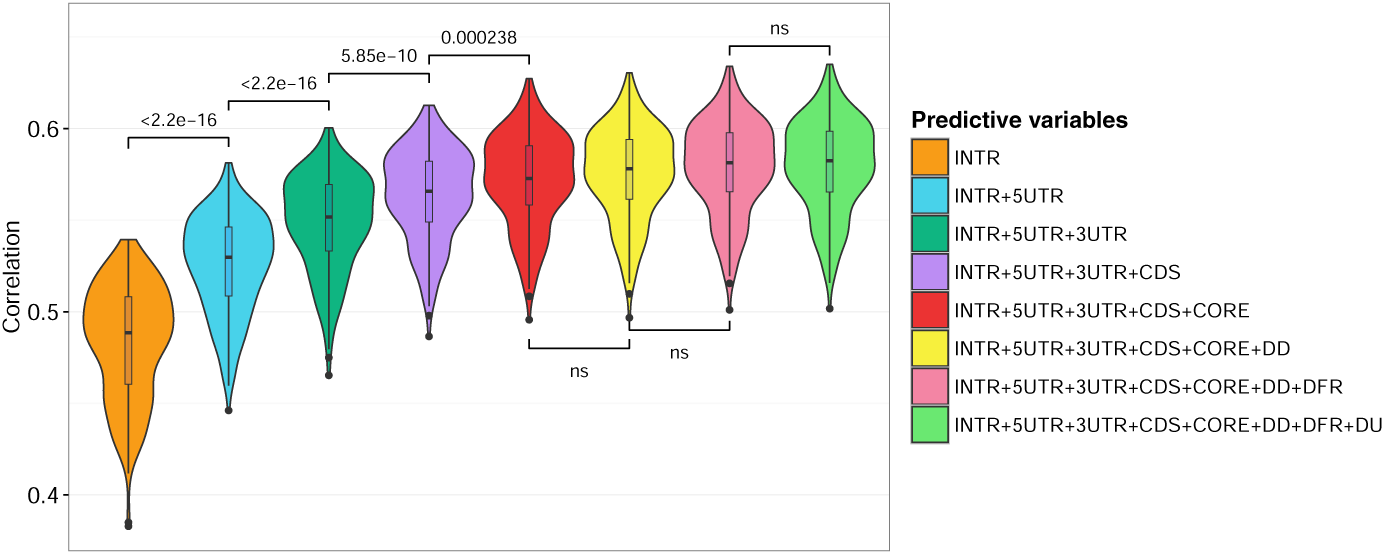
Contribution of additional genomic regions. Genomic regions were ranked according to their contribution in predicting gene expression. First, all regions were tested separately. Introns yielded the highest Spearman correlation between observed and predicted expressions and was selected as the ‘first’ seed region. Second, each region not already in the model was added separately. 5UTR in association with introns yielded the best correlation and was therefore selected as the ‘second’ region. Third, the procedure was repeated till all regions were included in the model. The contribution of each region is then visualized starting from the most important (left) to the less important (right). Note that the distance between the second TSS and the first ATG is > 2000 bp for only 189 genes implying that 5UTR and DD regions overlap. The correlations computed at each steps are indicated in Supplementary Table S1. ns, non significant.

Because RNA-associated regions (introns, UTRs, CDSs) had greater contribution to the prediction accuracy compared to DNA regions (promoters, DFR), we compared the accuracy of our model in predicting gene vs. transcript expression. We retrieved the normalized results for gene expression (RNAseqV2 rsem.genes.normalized_results) and the matched normalized expression signal of individual isoforms (RNAseqV2 rsem.isoforms.normalized_results) for 225 TCGA samples. Accordingly, we generated a set a predictive variables specific to each isoform (see Material and Methods). We found that models built on isoforms are less accurate than models built on genes (median r = 0.35, Supplementary Figure S5 and Table S2). This results is likely due to the fact that reconstructing and quantifying full-length mRNA transcripts is a difficult task, and no satisfying solution exists for now [51]. Consequently isoform as opposed to gene expression is more difficult to measure and thus to predict.

### Selecting DNA features related to gene expression

We sought the main DNA features related to gene expression. The complete model built on all 8 regions (160 variables) selected ~ 129 predictive variables per sample. We used the stability selection algorithm developed by Meinshausen *et al.* [52] to identify the variables that are consistently selected after data subsampling (see Materials and Methods for a complete description of the procedure). This procedure selected ~ 16 variables per sample. The barplot in Figure 5(a) shows, for each variable, the proportion of samples in which the variable is selected with high consistency (> 70% of the subsets).

**Figure 5.**
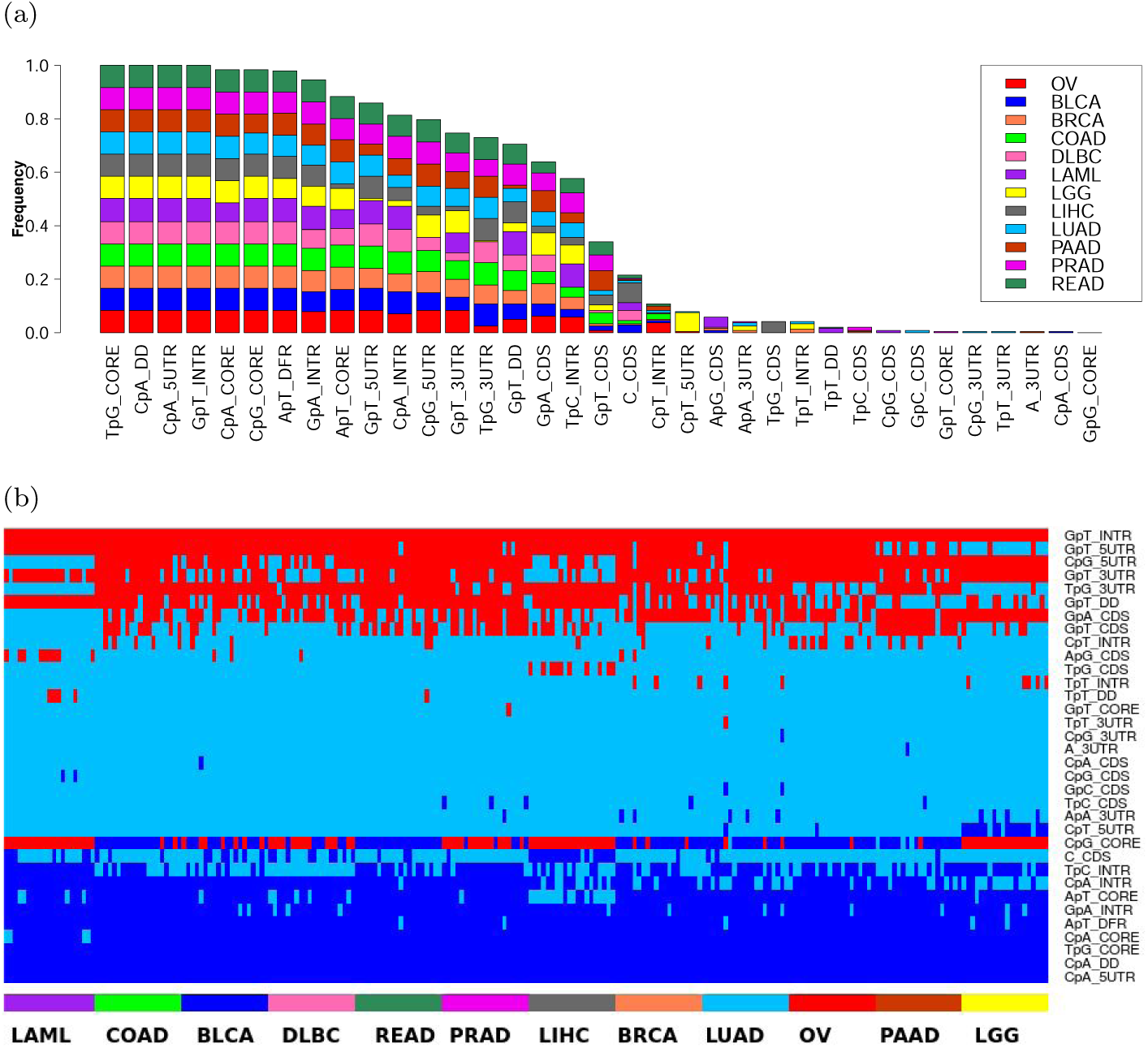
**(a) Consistently selected variables among 12 types of cancer.** For each variable, the fraction of samples in which the variable is considered as stable (i. e. selected in more than 70% of the subsets after subsampling) is shown. Each color refers to a specific type of cancer. Only variables consistently selected in at least one sample are shown (out of the 160 variables). See Materials and Methods for stable variable selection procedure and cancer acronyms. **(b) Biological effect of the stable variables**. For each of the 241 samples (columns), a linear model was fitted using the variables (rows) stable for this sample only. The sign of the contribution of each variable in each sample is represented as follows: red for positive contribution, dark blue for negative contribution and sky blue refers to variables not selected (i.e. not stably selected for the considered sample). Only the variables stable in at least one sample are represented. Cancers and samples from the same cancer types are ranked by decreasing mean error of the linear model.

We next determined whether stable variables exert a positive (activating) or a negative (inhibiting) effect on gene expression. For each sample, we fitted a linear regression model predicting gene expression using only the standardized variables that are stable for this sample. The activating/inhibiting effect of a variable is then indicated by the sign of its regression coefficient: < 0 for a negative effect and > 0 for a positive effect. The outcome of these analyses for all variables and all samples is shown Figure 5(b). With the noticeable exception of CpG in the core promoter, all stable variables had an invariable positive (e.g. GpT in introns) or negative (e.g. CpA in DD and in 5UTR) contribution in gene expression prediction in all samples. In contrast, CpG in the core promoter had an alternating effect being positive in LAML and LGG for instance while negative in READ. It is also the only variable with a regression coefficient close to 0 (absolute value of median = 0.1, see Supplementary Figure S6), providing a partial explanation for the observed changes. As CpG methylation inhibits gene expression [44], we also investigated potential differences in core promoter methylation in LAML (positive contribution of CpG_CORE) and READ (negative contribution of CpG_CORE). We used the Illumina Infinium Human DNA Methylation 450 made available by TCGA and focused on the estimated methylation level (beta values) of the sites intersecting with the core promoter. We noticed that core promoters in LAML were overall more methylated (median = 0.85) than in READ (median = 0.69, wilcoxon test p-value < 2.2e-16), opposite to the sign of CpG coefficient in LAML (positive contribution of CpG_CORE) and READ (negative contribution of CpG_CORE). This argued against a contribution of methylation in the alternating effect of CpG_CORE.

In order to characterize well predicted genes, we used a regression tree [53] to classify genes according to the prediction accuracy of our model (i.e. absolute error). The nucleotide and dinucleotide compositions of the various considered regions were used as classifiers. This approach identified groups of genes with similar (di)nucleotide composition in the regulatory regions considered and for which our model showed similar accuracy (Supplemental Figure S7). Implicitly, it identified the variables associated with a better or a poorer prediction. We applied this approach to the 241 linear models. The number of groups built by a regression tree differs from one sample to another (average number = 14). The resulting 3,680 groups can be visualized in the heatmap depicted in Figure 6, wherein each column represents a sample and each line corresponds to a group of genes identified by a regression tree. This analysis showed that our model is not equally accurate in predicting the expression of all genes but mainly fits certain classes of genes (bottom rows of the heatmap, Figure 6) with specific genomic features (Supplementary Figure S7). Note that the groups well predicted in all cancers likely correspond to ubiquitously expressed (ie. housekeeping) genes (Supplementary Figure S7(a) and Supplementary Table S3). In contrast, some groups were well predicted in only certain cancers and were associated to specific biological function (compare for instance white groups of PAAD and LGG vs. LAML, DBLC and LIHC in Figure 6, Supplementary Figure S7(b) and Supplementary Table S4).

**Figure 6.**
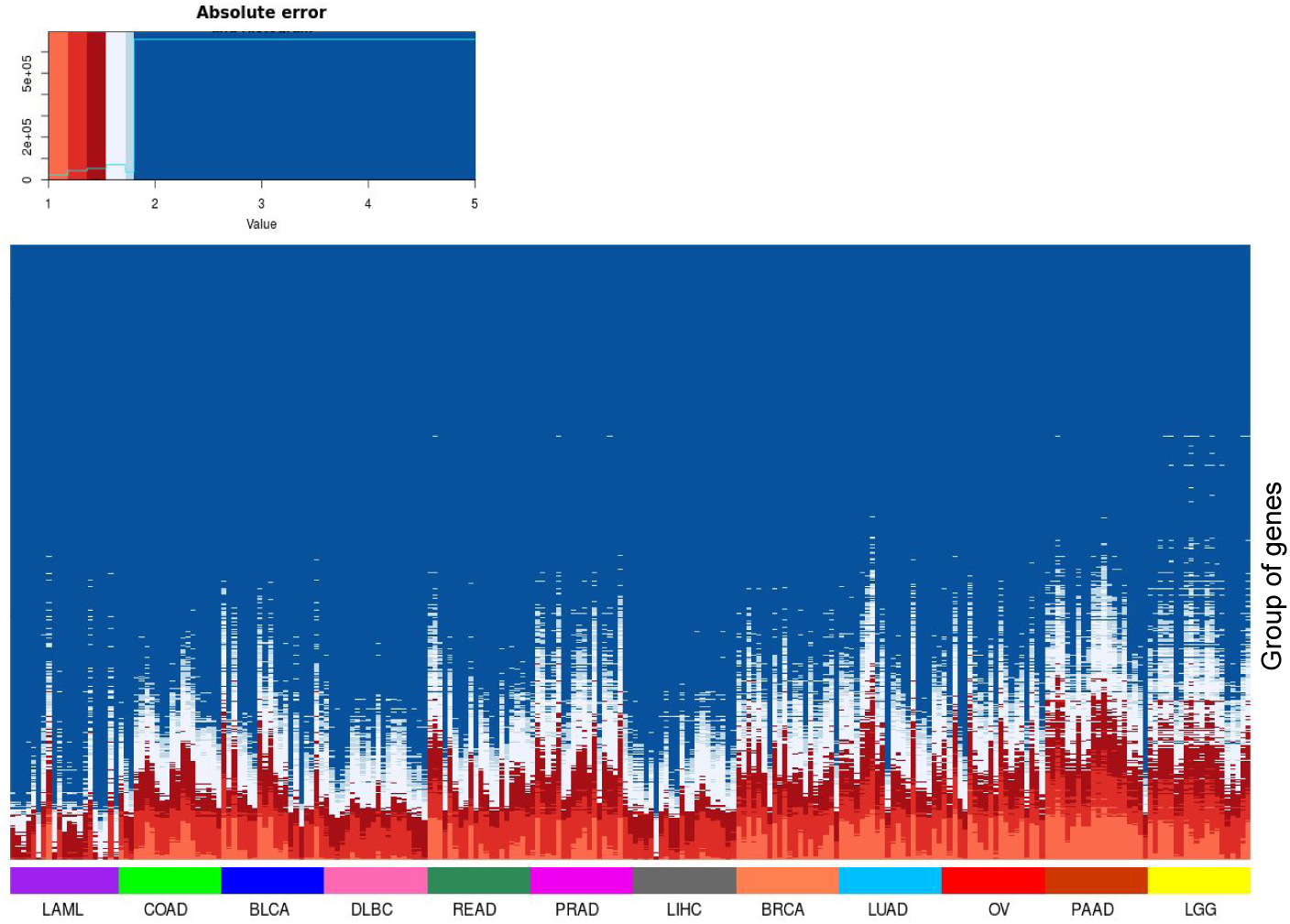
Gene classification according to prediction accuracy. Columns represent the various samples gathered by cancer type. Samples from the same cancer type are ranked by decreasing mean squared prediction error. Lines represent the 3,680 groups of gene obtained with the regression trees (one tree for each of the 241 samples) ranked by decreasing mean squared prediction error. Groups gathering the top 25% well predicted genes (error <~1.77) are indicated in red and light blue.

### Relationships between selected nucleotide composition and genome architecture

We probed the regulatory activities of the selected regions. We first noticed that several regions considered, in particular introns and DFRs, contain enhancers mapped by the FANTOM consortium [7] (Supplementary Table S5). Enhancers and promoters share several similarities including the binding of TFs through discrete motifs [1], that can be affected by genomic variations [54]. We then used the v6p GTEx release to compute the average frequencies of *cis* expression quantitative trait loci (*cis*-eQTLs) present in the considered genomic regions and directly linked to their host genes (Supplementary Table S6). Introns contained the smallest number of eQTLs (10 times less than any other regions), indicating that these sequences, as opposed to DFRs and core promoters for instance (Supplementary Table S6), harbor few specific motifs, in particular TF binding motifs, that can be affected by single nucleotide variations. It was therefore unlikely that the effect of introns was solely due to the presence of intronic enhancers. It rather unveiled the existence of another layer of regulation, which is not mediated by specific motifs but presumably involves larger DNA regions. Note that, because introns represent the most important region contributing to our model, *cis*-eQTL frequencies also confirmed that our model is more efficient in evaluating the contributions of nucleotide environment than that of short motifs, as already suggested by the secondary contribution of TF motifs and DNA shapes (Figure 2(d)).

We then asked whether the groups of genes identified by the regression trees (Figure 6) correspond to specific TADs. Genes within the same TAD tend to be coordinately expressed [55, 56]. TADs with similar chromatin states tend to associate to form two genomic compartments called A and B: A contains transcriptionally active regions while B corresponds to transcriptionally inactive regions [57]. The driving forces behind this compartmentalization and the transitions between compartments observed in different cell types are not fully understood, but chromatin composition and transcription are supposed to play key roles [5]. We reasoned that the groups of genes identified by the regression trees reflect gene associations within TADs and beyond assignment of TADs to compartments A or B. This rationale implied first that TADs can be distinguished according to the nucleotide composition of their resident genes. We used the 373 TADs containing more than 10 genes mapped in IMR90 cells [6]. For each TAD, we compared the nucleotide compositions of the embedded genes and the nucleotide compositions of all other genes using a Kolmogorov-Smirnov test. We used a Benjamini-Hochberg multiple testing correction to control the False Discovery Rate (FDR), which was fixed at 0.05. We found that 324 TADs out of 373 (~87%) are characterized by at least one specific nucleotide signature (Figure 7(a)). In addition, our results clearly showed the existence of distinct classes of TADs related to GC content (GC-rich, GC-poor and intermediate GC content) (Figure 7(a)). This is in line with the results of Jabbari and Bernardi, who showed that the distribution of GCs along the genome (i.e. isochores) can help define TADs [58].

We next considered the 967 groups of genes defined in Figure 6 whose expression is accurately predicted by our model (i.e. groups with mean error < mean error of the 1st quartile). We thus focused our analyses on genes for which we did learn some regulatory features. We evaluated the enrichment for specific TADs in each group (considering only TADs containing more than 10 genes) using an hypergeometric test (Figure 7(b)). We found that 60% of these groups were enriched for at least one TAD (p-value < 0.05). Hence, several groups of genes identified by the regression trees (Figure 6) do correspond to specific TADs (Figure 7(b)). Overall our results are in agreement with the idea that TADs regroup genes according to their nucleotide compositions.

**Figure 7.**
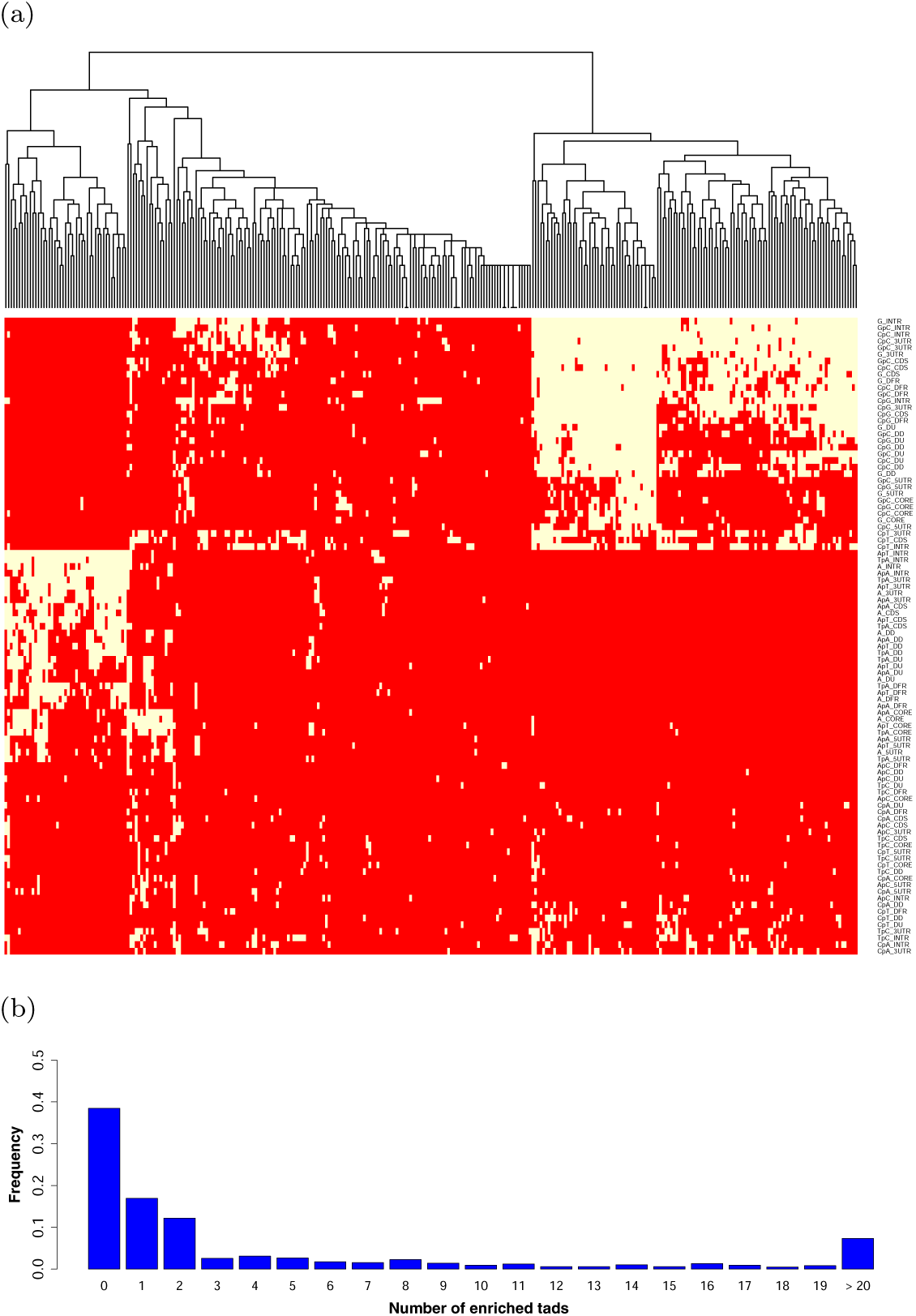
**(a) Nucleotide compositions of resident genes distinguish TADs.** For each TAD and for each region considered, the percentage of each nucleotide and dinucleotide associated to the embedded genes were compared to that of all other genes using a Kolmogorov-Smirnov test. Red indicates FDR-corrected p-value ≥ 0.05 and yellow FDR-corrected p-value < 0.05. TAD clustering was made using this binary information. Only TADs with at least one p-value < 0.05 are shown (i.e. 87% of the TADs containing at least 10 genes**) (b) TAD enrichment within groups of genes whose expression is accurately predicted by our model.** The enrichment for each TAD (containing more than 10 genes) in each gene group accurately predicted by our model (i.e. groups with mean error < mean errors of the 1st quartile) was evaluated using an hypergeometric test. The fraction of groups with enriched TADs (p-value < 0.05) is represented.

## DISCUSSION

In this study, we challenged the hypothesis that DNA sequence contains information able to explain gene expression [31, 32, 33, 34, 35]. We built a global regression model to predict, in any given sample, the expression of the different genes using only nucleotide compositions as predictive variables. Overall our model provided a framework to study gene regulation, in particular the influence of regulatory regions and their associated nucleotide composition.

A collateral and surprising result of our study is the limited biological information brought by linear models built on experimental data (ChIP-seq and DNaseI-seq) [12, 30]. The similar accuracy of these models on real and randomly permuted data indicated that, though the experimental data are biologically relevant, their interpretation through a linear model is not straightforward. An interesting perspective would be to devise a strategy to infer TF combinations from experimental data without being influenced by the opening of the chromatin.

The accuracy of our model confirmed that DNA sequence *per se* and basic information like dinucleotide frequencies have very high predictive power. It remains to determine the exact nature of these sequence-level instructions. Interestingly, nucleotide environment contributes to prediction of TF binding sites and motifs bound by a TF have a unique sequence environment that resembles the motif itself [47]. Hence, the potential of the nucleotide content to predict gene expression may be related to the presence of regulatory motifs and TFBSs. However, we showed that the gene body (introns, CDS and UTRs), as opposed to sequences located upstream (promoter) or downstream (DFR), had the most significant contribution in our model. Moreover, *cis*-eQTL frequencies argue against the presence of short regulatory motifs notably in introns, suggesting the existence of another layer of regulation that implicates large DNA regions.

We indeed provided evidence that the contribution of nucleotide composition in predicting gene expression might be linked to co-regulations associated with genome 3D architecture. We specifically showed that TADs regroup genes according to their nucleotide composition. In line with our results, Kornyshev *et al.* have shown that physical attractive forces between DNA fragments rely on sequence homology [59]. Paralog genes, which are generated by tandem duplication and therefore have similar nucleotide composition, are co-regulated according to TADs [60]. Strikingly, Singh *et al.* developed deep learning models able to predict enhancer-promoter interactions based on sequence-based features only [61]. These results are reminiscent of the sequence-encoded ‘‘enhancer-corepromoter specificity” observed in Drosophila [62]. The same rationale might be envisaged for gene interactions within TADs as enhancers prefer to activate promoters resembling those of their parent genes [62, 63]. These results together with our findings suggest that our genome encodes sequence-level instructions that help determine genomic interactions. Although the sequence instructions encoded by genomic DNA are - almost - identical in all cell types of an individual, their usage must however be cell-type specific to allow the proper A/B compartimentalization of TADs and ultimately the diversity of cell types and functions. At this stage, the mechanisms driving this cell-type specific selection of nucleotide compositions remain to be characterized.

## ACKNOWLEDGEMENT

We thank Mohamed Elati, Mathieu Lajoie, Anthony Mathelier and Cédric Notredame for insightful discussions and suggestions. We also thank Yue Li, Zhaolei Zhang, Florian Schmidt and Marcel H. Schulz for sharing data. We are indebted to the researchers around the globe who generated experimental data and made them freely available. C-H.L. is grateful to Marc Piechaczyk, Edouard Bertrand, Anthony Mathelier and Wyeth W. Wasserman for continued support.

